# The role of pregnancy-specific glycoproteins on trophoblast motility in three-dimensional gelatin hydrogels

**DOI:** 10.1101/2020.09.25.314195

**Authors:** Samantha G. Zambuto, Shemona Rattila, Gabriela Dveksler, Brendan A.C. Harley

## Abstract

Trophoblast invasion is a complex biological process necessary for establishment of pregnancy; however, much remains unknown regarding what signaling factors coordinate the extent of invasion. Pregnancy-specific glycoproteins (PSGs) are some of the most abundant circulating trophoblastic proteins in maternal blood during human pregnancy, with maternal serum concentrations rising to as high as 200-400 μg/mL at term. Here, we employ three-dimensional (3D) trophoblast motility assays consisting of trophoblast spheroids encapsulated in 3D gelatin hydrogels to quantify trophoblast outgrowth area, viability, and cytotoxicity in the presence of PSG1 and PSG9 as well as epidermal growth factor and Nodal. We show PSG9 reduces trophoblast motility whereas PSG1 increases motility. Further, we assess bulk nascent protein production by encapsulated spheroids to highlight the potential of this approach to assess trophoblast response (motility, remodeling) to soluble factors and extracellular matrix cues. Such models provide an important platform to develop a deeper understanding of early pregnancy.

**Graphical Abstract:** 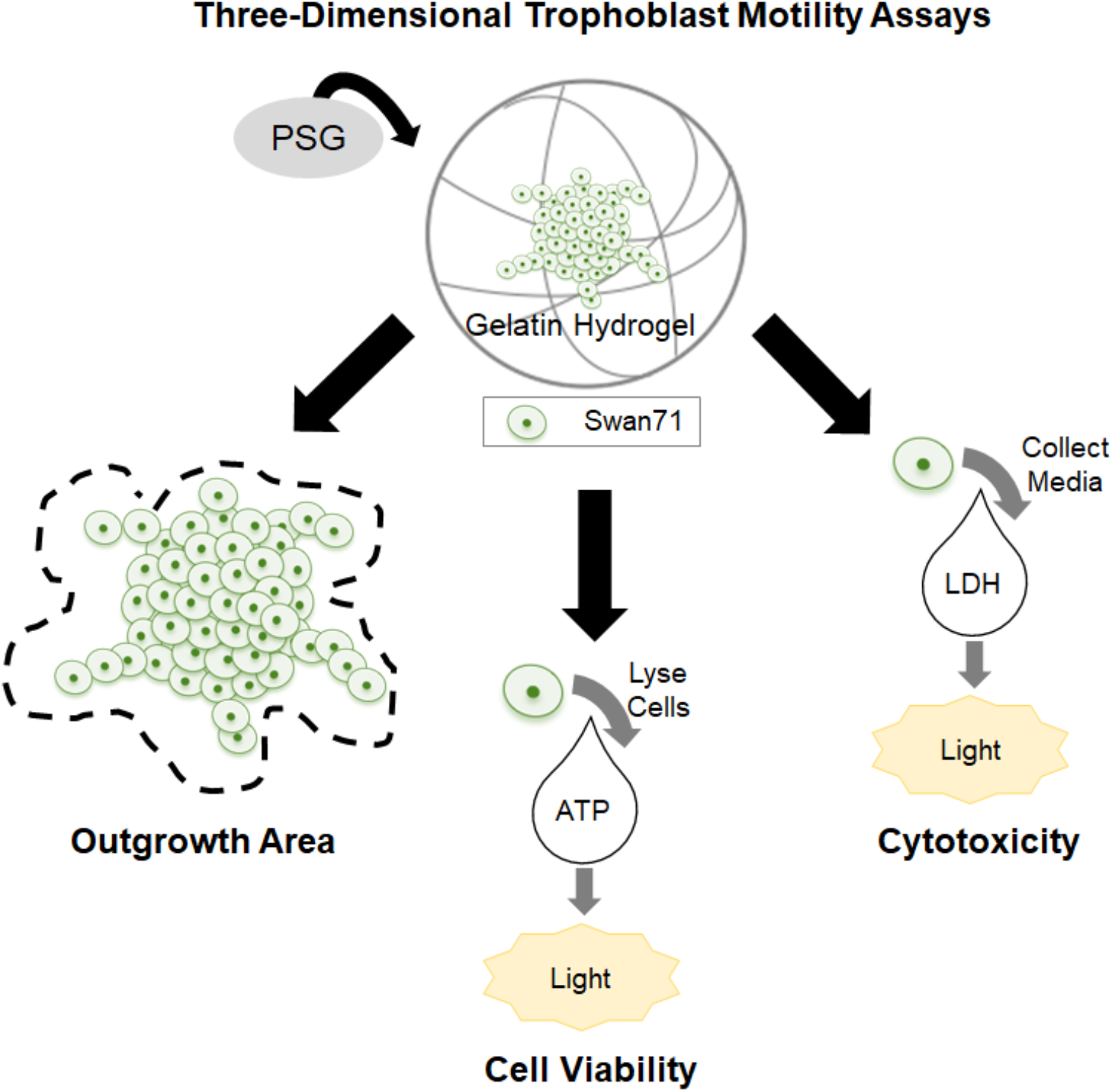

## 1. Introduction

Pregnancy is an intricate biological process that involves a complex molecular dialogue between cells from the endometrium and trophoblast cells from the invading blastocyst (Cha et al., 2012; Norwitz et al., 2001); however, what orchestrates this cellular crosstalk remains poorly understood. Blastocyst implantation into the endometrium is thought to occur across three phases (Cha et al., 2012; Norwitz et al., 2001; Su and Fazleabas, 2015). Apposition is the initial, unstable connection between the endometrium and the blastocyst (Cha et al., 2012; Norwitz et al., 2001; Su and Fazleabas, 2015). Adhesion establishes a more stable, physical connection between the endometrium and blastocyst (Cha et al., 2012; Norwitz et al., 2001; Su and Fazleabas, 2015). Finally, invasion occurs when trophoblasts breach the endometrial epithelium and embed the blastocyst within the endometrial stromal tissue (Cha et al., 2012; Norwitz et al., 2001; Su and Fazleabas, 2015). Ultimately, implantation is modulated by molecular signaling between cells from a hormonally-primed endometrium and trophoblast cells from a mature blastocyst (Cha et al., 2012; Norwitz et al., 2001; Su and Fazleabas, 2015). Although researchers have highlighted the importance of biological stimuli such as ovarian hormones estrogen and progesterone that modulate endometrial decidualization for successful implantation, much remains unknown regarding other factors expressed by maternal cells and trophoblast that coordinate the extent of invasion (Cha et al., 2012; Norwitz et al., 2001; Su and Fazleabas, 2015). Developing a deeper understanding of what biological cues influence invasion would provide clinical insights that may improve our ability to understand, prevent, and treat fertility disorders and early pregnancy loss.

Numerous challenges exist regarding studying early implantation in humans. Notably, this process occurs shortly after conception: the blastocyst is embedded within endometrial tissue by approximately 10 days post-conception which provides a significant challenge to directly studying this process within humans (Cha et al., 2012; Norwitz et al., 2001; Su and Fazleabas, 2015). As such, initial adhesion of the blastocyst to the endometrium has never been observed in humans and much of what we know regarding early pregnancy is inferred from rare histological specimens or from animal studies (Cha et al., 2012; Norwitz et al., 2001; Su and Fazleabas, 2015). Nevertheless, significant differences between human pregnancy and placentation compared to rodents and non-human primates calls into question the relevance of using such models to study mechanisms of early implantation in humans (Cha et al., 2012; Norwitz et al., 2001; Su and Fazleabas, 2015).

Three-dimensional (3D) models, such as organoid models and tissue engineered platforms, offer the unique opportunity to elucidate mechanistic processes associated with human implantation that are currently impossible to study in the body with current state of the art technologies. Unlike animal models which are complex and costly, these *in vitro* platforms offer controllability and scalability while providing an avenue to incorporate increasingly complex culture environments (e.g., cells, extracellular matrix, biomolecules) relevant to what occurs in the body (Cook et al., 2017; Valdez et al., 2017; Zambuto et al., 2019a). Here, we use methacrylamide-functionalized gelatin (GelMA) hydrogels to investigate the role of pregnancy-specific glycoproteins (PSGs) on trophoblast motility. Gelatin is an attractive platform for these studies because it is denatured collagen that retains relevant arginine-glycine-aspartic acid (RGD) cell binding motifs and matrix metalloproteinase-sensitive (MMP) degradation sites that ultimately allow for cellular adhesion and matrix remodeling which are critical processes in invasion (Pedron and Harley, 2013; Zambuto et al., 2019b). The addition of methacrylamide groups to gelatin’s backbone allows for photopolymerization, which renders GelMA hydrogels stable under physiological temperatures and relatively homogeneous in terms of structure (Pedron and Harley, 2013; Zambuto et al., 2019b). Such models have previously been employed for quantifying trophoblast and cancer cell invasion in 3D (Chen et al., 2018c; Zambuto et al., 2019b, 2020). Here, we adapted the technique to track cell migration away from cell spheroids embedded within the hydrogel (Chen et al., 2018a; Chen et al., 2018b; Pedron et al., 2019) to investigate the pattern of motility by Swan71 trophoblast cells in the presence of soluble factors present at the maternal-fetal interface. In contrast to 2D assays such as wound healing assays and Boyden chamber assays, these 3D motility strategies can replicate aspects of the *in vivo* environment, including tissue stiffness, 3D tissue architecture, and matrix composition.

In this work, we describe an approach to compare trophoblast motility patterns in response to soluble pregnancy-specific glycoproteins (PSGs). PSGs are some of the most abundant circulating trophoblastic proteins in maternal blood during human pregnancy (Moore and Dveksler, 2014; Rattila et al., 2019; Sorensen, 1984; Wurz et al., 1981). Over the course of pregnancy, the maternal serum level of PSGs increases to approximately 200-400 μg/mL at term (Moore and Dveksler, 2014). There are 10 coding *PSG* genes in humans, *PSG1-PSG9* and *PSG11*, and one non-coding pseudogene *PSG10* (Moore and Dveksler, 2014; Rattila et al., 2019). At the end of the first trimester, *PSG1* and *PSG3* account for the bulk majority of expression and by term, *PSG1, PSG3, PSG4, PSG5*, and *PSG6* are equally expressed with low expression of the other *PSGs* (Moore and Dveksler, 2014). PSGs are expressed by extravillous trophoblasts and syncytiotrophoblasts (Rattila et al., 2019; Zhou et al., 1997). Lower serum concentrations of PSGs have been associated with pregnancy disorders and pathologies, such as fetal growth restriction and preeclampsia (Moore and Dveksler, 2014; Rattila et al., 2019). For example, reduced serum concentrations of PSG1 were reported in African American women diagnosed with early- and late-onset preeclampsia but only for male fetuses (Rattila et al., 2019). Upregulation of PSG9 expression was found in colorectal carcinogenesis (Moore and Dveksler, 2014); however, much remains unknown regarding the role of PSG9 on trophoblast invasion. This study quantified the differences in trophoblast outgrowth area, fold change in outgrowth area, viability, and cytotoxicity after addition of PSG1 or PSG9 to the cell media. We report differences in outgrowth area and viability/cytotoxicity based on the addition of certain factors and investigate whether encapsulated trophoblast cells exhibit hallmarks of extracellular matrix remodeling within the hydrogels.

## 2. Results

### 2.1. Characterization of Swan71 Trophoblast Spheroid Motility Assays

A library of Swan71 spheroids was created with diameters that fall within the regime of invading blastocysts (Shapiro et al., 2008). Spheroids fabricated from 2,000, 4,000, or 6,000 cells/spheroid ranged in size from rom 360.4 ± 10.3 μm to 535.0 ± 11.1 μm in diameter and 0.11 ± 0.01 to 0.21 ± 0.03 mm^2^ in projected area (Figure 1A-C). All spheroid densities were significantly different from each other in diameter (n=5 per condition; One-way ANOVA; *p* < 0.0001) and area (n=5; One-Way ANOVA; 2,000 to 4,000 cells, *p*=0.0016; 2,000 to 6,000, *p*=4.14×10^−6^; 4,000 to 6,000 *p*=0.0037). Spheroids of 4,000 cells were selected for all subsequent studies and were encapsulated in GelMA hydrogels for up to 3 days (Figure 1D). By day 3 of culture, trophoblast cells were observed to migrate into the surrounding hydrogel matrix away from the initial spheroid core (Figure 1E).

**Figure 1.**
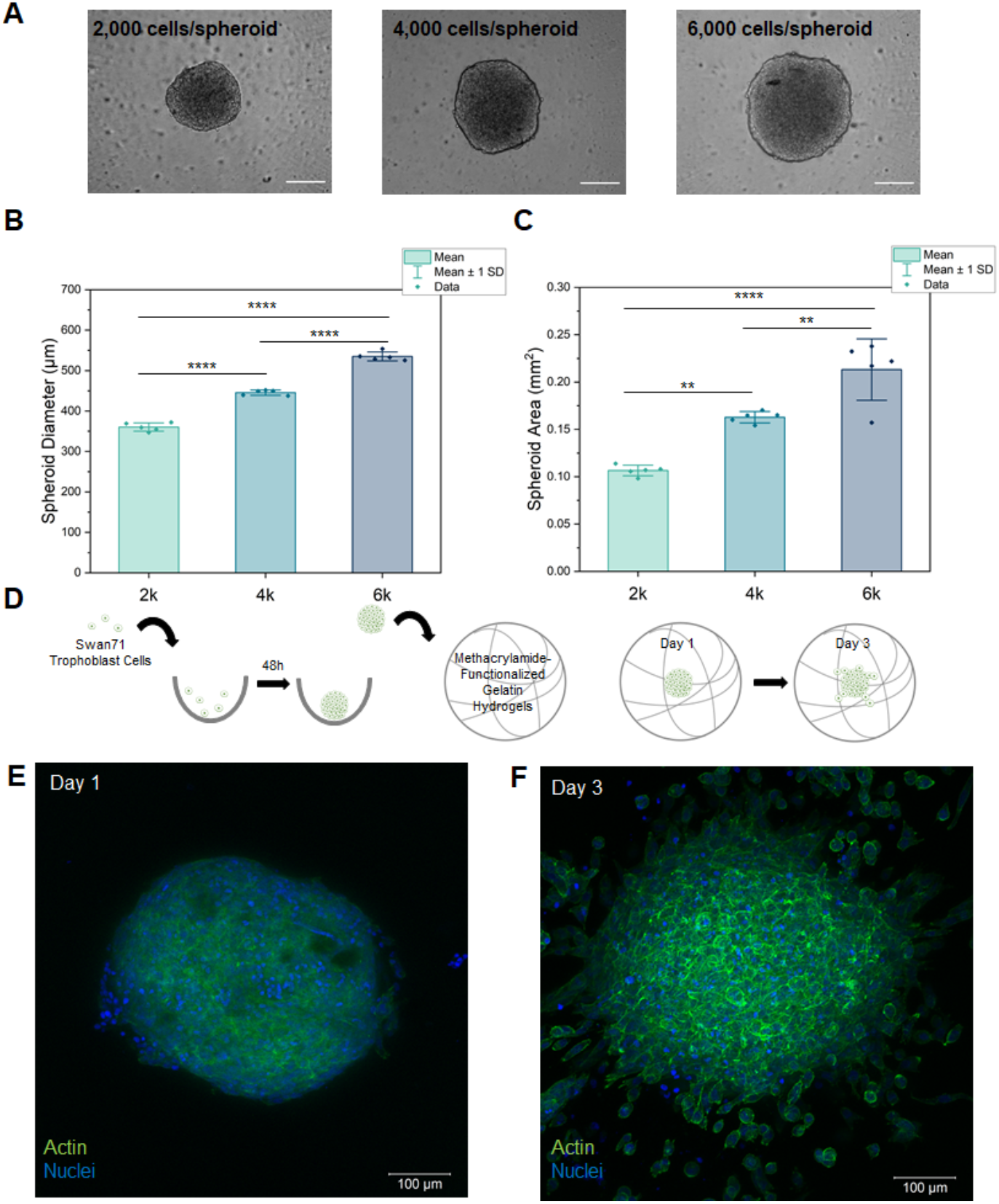
Three-Dimensional Swan71 Trophoblast Spheroid Assays. (A) Representative bright field images of Swan71 spheroids (2,000, 4,000, and 6,000 cells/spheroid). Scale bar, 200 μm. (B) Average spheroid diameter in round bottom plates. Data are displayed as individual data points. Bar represents mean ± standard deviation. One-way ANOVA with Tukey post hoc analysis, **** p < 0.0001 (n=5 technical replicates per group). (C) Average spheroid area in round bottom plates. Data are displayed as individual data points. Bar represents mean ± standard deviation. One-way ANOVA with Tukey post hoc analysis, ** p < 0.01, **** p < 0.0001 (n=5 technical replicates per group). (D) Schematic of experimental procedure. (E) Representative maximum intensity projection of encapsulated Swan71 spheroid on day 1 of culture. Scale bar, 100 μm. (F) Representative maximum intensity projection of encapsulated Swan71 spheroid on day 3 of culture. Scale bar, 100 μm.

### 2.2. Epidermal Growth Factor Increases Trophoblast Motility and Viability Compared to Control and to Nodal

To demonstrate that soluble biomolecules significantly alter baseline motility patterns of Swan71 trophoblasts, we encapsulated Swan71 spheroids for 3 days in the presence of the known invasion promoter epidermal growth factor (EGF; 5 ng/mL) and known invasion inhibitor Nodal (250 ng/mL) (Kuo et al., 2016; Nadeem et al., 2011; Staun-Ram et al., 2004). Over 3 days of culture, trophoblast cells migrated into the surrounding hydrogel matrix and we detected quantifiable differences in migration patterns and cell viability (Figure 2). By day 2, cells cultured in medium containing EGF had significantly higher outgrowth area (Figure 2B; Kruskal-Wallis ANOVA, n=6-7 per condition) compared to cells cultured in control medium (*p*=0.022) or medium containing recombinant nodal (*p*=1.97×10^−3^). By day 3, we observed robust differences (One-way ANOVA, n=6-7) in outgrowth area in samples cultured with EGF compared to control and recombinant Nodal (*p*<0.0001). Differences in fold change in outgrowth area (Figure 2C) of samples compared to initial spheroid area on day 0 were discernable by day 1 for EGF samples compared to nodal samples (One-way ANOVA, n=6-7, *p*=0.048). By days 2 and 3, EGF samples had significantly higher fold change compared to control and nodal samples (Day 2: Welch’s Heteroscedastic F Test with Trimmed Means and Winsorized Variances, n=6-7, EGF Control *p*=0.006, EGF Nodal *p*=0.004; Day 3: One-way ANOVA, n=6-7, EGF Control *p*=4.22×10^−5^, EGF Nodal *p*=1.76×10^−5^). While cell viability (Figure 2D) was increased for samples cultured with EGF compared to control samples (Welch’s ANOVA, n=6-7, *p*=0.033) there were no statistically significant differences in cytotoxicity (Figure 1E) between the three groups by day 3 (One-way ANOVA, n=5-7). Collectively, these data demonstrate that 3D spheroid motility assays provide an engineering platform that can be used to detect shifts in motility and viability/cytotoxicity patterns in response to soluble biomolecules.

**Figure 2.**
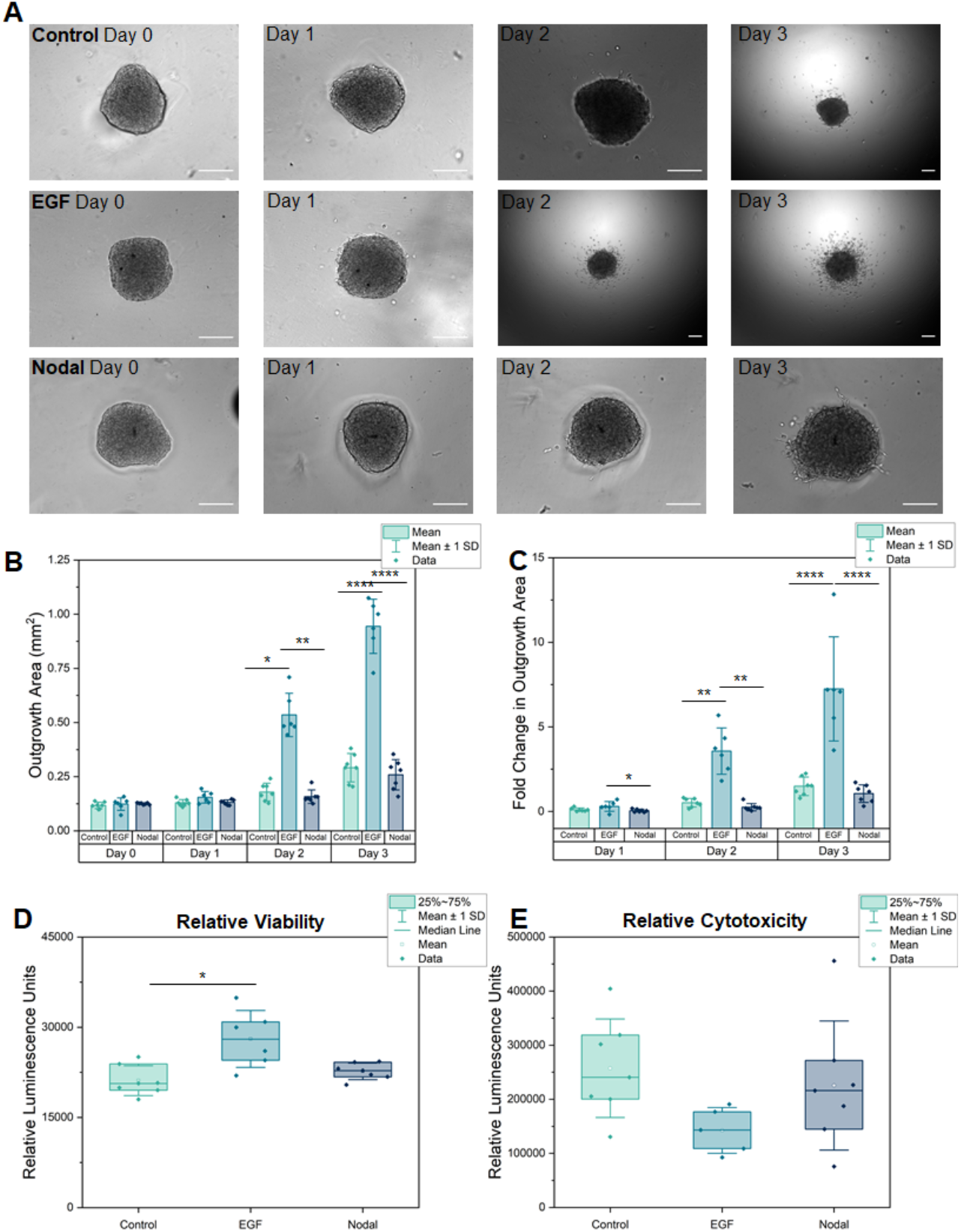
Invasion Pattern Analysis After the Addition of Exogenous Growth Factors Epidermal Growth Factor (EGF) and Nodal to Spheroids Encapsulated in Methacrylamide-Functionalized Gelatin (GelMA) Hydrogels. (A) Representative bright field images of 4,000 cell spheroids encapsulated in GelMA hydrogels from day 0 (seeding) to day 3 for control and after addition of 5 ng/mL EGF or 250 ng/mL Nodal. Scale bar, 200 μm. (B) Average spheroid outgrowth area in GelMA hydrogels over 3 days. Data are displayed as individual data points. Bar represents mean ± standard deviation. Data were analyzed on each day between groups (Control n=7 technical replicates, EGF n=6 technical replicates, Nodal n=7 technical replicates). Day 0: One-way ANOVA with Tukey post hoc analysis, *** p < 0.001, **** p < 0.0001. Day 1: One-way ANOVA with Tukey post hoc analysis. Day 2: Kruskal-Wallis with Dunn’s post hoc analysis, * p < 0.05, ** p < 0.01. Day 3: One-way ANOVA, **** p < 0.0001. (C) Spheroid outgrowth area fold change compared to initial area (day 0) over 3 days. Data are displayed as individual data points. Bar represents mean ± standard deviation. Data were analyzed on each day between groups (Control n=7 technical replicates, EGF n=6 technical replicates, Nodal n=7 technical replicates). Day 1: One-way ANOVA with Tukey post hoc analysis, * p < 0.05. Day 2: Welch’s heteroscedastic F Test with Trimmed Means and Winsorized Variances with Games-Howell post hoc analysis, ** p < 0.01. Day 3: One-way ANOVA with Tukey post hoc analysis, **** p < 0.0001. (D) Relative viability of encapsulated spheroids at day 3 from CellTiter-Glo® 3D Viability Assay. Data are presented as individual data points overlaying box plots with the median denoted by a line, mean denoted by a square, and whiskers represent the mean ± standard deviation. Welch’s ANOVA with Games-Howell post hoc analysis, * p < 0.05, Control n=7 technical replicates, EGF n=6 technical replicates, Nodal n=7 technical replicates. (E) Relative cytotoxicity of encapsulated spheroids at day 3 from measured from lactate dehydrogenase release via LDH-Glo® Cytotoxicity Assay. Data are presented as individual data points overlaying box plots with the median denoted by a line, mean denoted by a square, and whiskers represent the mean ± standard deviation. One-way ANOVA with Tukey post hoc analysis, Control n=7 technical replicates, EGF n=5 technical replicates, Nodal n=7 technical replicates.

### 2.3. The Addition of Soluble PSG9-Fc Decreases Trophoblast Motility

We subsequently quantified changes in trophoblast spheroid outgrowth area, viability, and cytotoxicity over 3 days of culture in response to 60 μg/mL soluble PSG9-Fc (Figure 3). By day 3 of culture, we observed a statistically significant decrease in outgrowth area (Figure 3B) in response to PSG9-Fc compared to both cell medium control and the Fc-treated control protein samples (One-way ANOVA, n=8 per condition, Control PSG9-Fc *p*=7.13×10^−6^, Fc-Control PSG9-Fc *p*=0.0066, Control Fc-Control *p*=0.020). The Fc-control was used as an additional control condition because PSG9-Fc has a Fc tag at the C-terminus; further, both PSG9-Fc and Fc-control were purified using the same procedure. Fold change (Figure 3C) in outgrowth area differed between control and PSG9-Fc samples on day 1 (Kruskal-Wallis ANOVA, n=8, *p*=0.019) and on day 3, fold change was significantly different (One-way ANOVA, n=8) in PSG9-Fc samples compared to control (*p*=9.60×10^−6^) and Fc-control samples (*p*=0.0022). Notably, day 3 cell viability (CellTiter-Glo® 3D Viability Assay; Figure 3D) did not differ between groups, though cytotoxicity (LDH-Glo® Cytotoxicity Assay; Figure 3E) was statistically significantly lower in Fc-control samples versus control media (One-way ANOVA, n=8, *p*=0.0018). Taken together, these data suggest that PSG9-Fc may have an inhibitory role on trophoblast invasion without influencing trophoblast viability or cytotoxicity.

**Figure 3.**
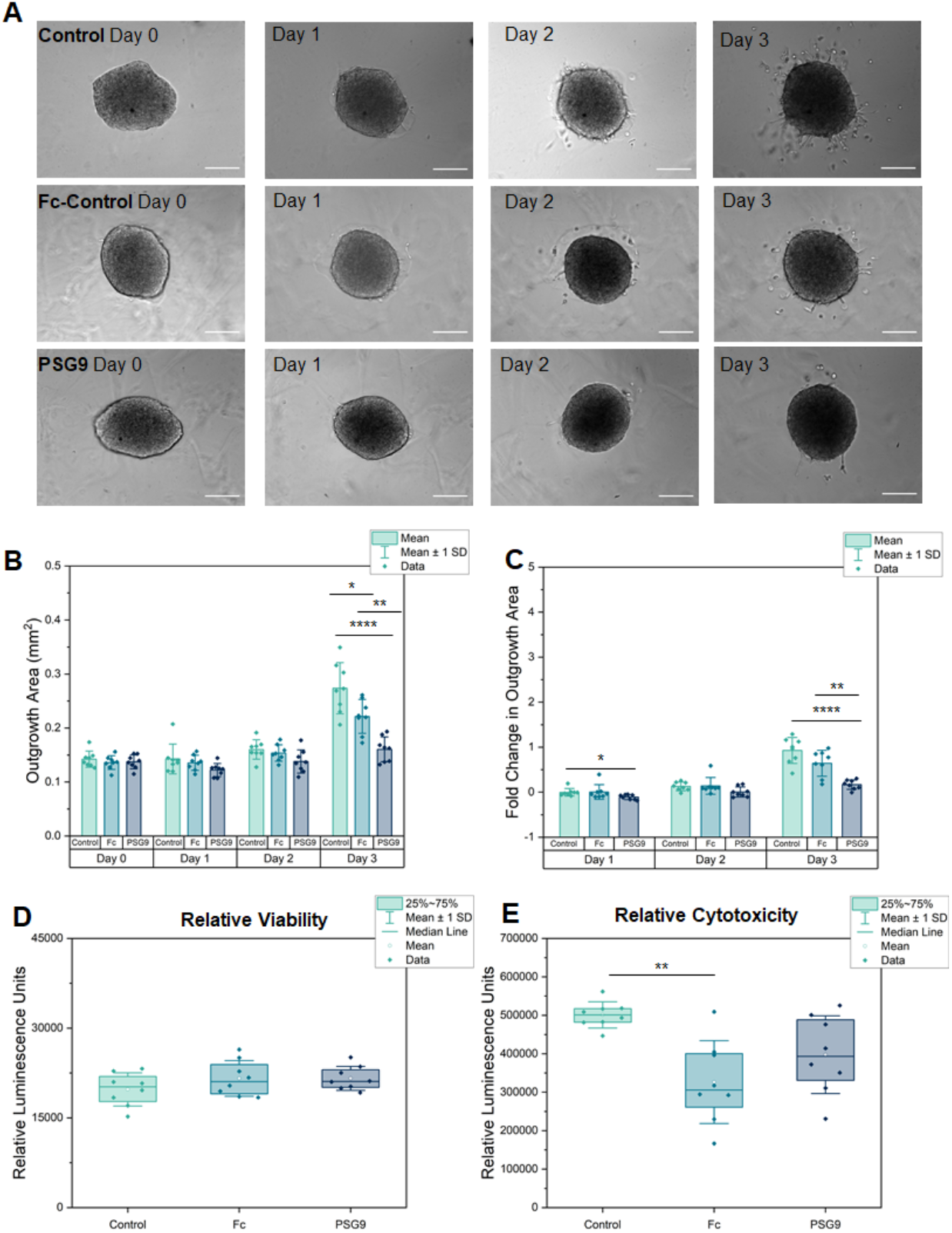
PSG9-Fc Inhibits Swan71 Spheroid Invasion in Methacrylamide-Functionalized Gelatin (GelMA) Hydrogels. (A) Representative bright field images of 4,000 cell spheroids encapsulated in GelMA hydrogels from day 0 (seeding) to day 3 for Control samples, Fc-Control samples (60 μg/ml), and PSG9-Fc samples (60 μg/ml). Scale bar, 200 μm. (B) Average spheroid outgrowth area in GelMA hydrogels over 3 days. Data are displayed as individual data points. Bar represents mean ± standard deviation. Data were analyzed on each day between groups (Control n=8 technical replicates, Fc-control n=8 technical replicates, PSG9-Fc n=8 technical replicates). Day 0: One-way ANOVA. Day 1: Kruskal-Wallis ANOVA. Day 2: One-way ANOVA. Day 3: One-way ANOVA with Tukey post hoc analysis, * p < 0.05, ** p < 0.01, **** p < 0.0001. (C) Spheroid outgrowth area fold change compared to initial area (day 0) over 3 days. Data are displayed as individual data points. Bar represents mean ± standard deviation. Data were analyzed on each day between groups (Control n=8 technical replicates, Fc-control n=8 technical replicates, PSG9-Fc n=8 technical replicates). Day 1: Kruskal-Wallis ANOVA with Dunn’s post hoc analysis, * p <0.05. Day 2: Kruskal-Wallis ANOVA. Day 3: One-way ANOVA with Tukey post hoc analysis, ** p < 0.01, **** p < 0.0001. (D) Relative viability of encapsulated spheroids at day 3 from CellTiter-Glo® 3D Viability Assay. Data are presented as individual data points overlaying box plots with the median denoted by a line, mean denoted by a square, and whiskers represent the mean ± standard deviation. One-way ANOVA. Control n=8 technical replicates, Fc-control n=8 technical replicates, PSG9-Fc n=8 technical replicates. (E) Relative cytotoxicity of encapsulated spheroids at day 3 measured from lactate dehydrogenase release via LDH-Glo® Cytotoxicity Assay. Data are presented as individual data points overlaying box plots with the median denoted by a line, mean denoted by a square, and whiskers represent the mean ± standard deviation. One-way ANOVA with Tukey post hoc analysis. Control n=8 technical replicates, Fc-control n=8 technical replicates, PSG9-Fc n=8 technical replicates; ** p < 0.01.

### 2.4. PSG1-Fc, but not PSG1-His, Increases Trophoblast Motility

Although reduced serum concentrations of PSG1 were previously found in women diagnosed with preeclampsia pregnant with a male, we examined the influence of PSG1 on trophoblast motility as insufficient trophoblast invasion has been associated with pre-eclampsia (Lala and Nandi, 2016; Rattila et al., 2019). Previously, Swan71 migration was shown to increase in response to immobilized PSG1 in two-dimensional *in vitro* studies, but no differences were observed in transwell invasion assays (Rattila et al., 2019). We quantified the role of soluble PSG1 on Swan71 trophoblast motility using a 3D motility assay. We compared patterns of migration and viability using two variants of PSG1 (Figure 4): PSG1-His and PSG1-Fc which differ based on their production and purification methods as previously described (Blois et al., 2014). By day 2 of culture, we observed higher outgrowth area (One-way ANOVA, n=5-6 spheroids per condition) in samples cultured with PSG1-Fc compared to control (*p*=0.026), Fc-control (*p*=0.029), and PSG1-His samples (*p*=8.42×10^−4^). On day 3 of culture, hydrogels cultured with PSG1-Fc had significantly higher (Welch’s ANOVA, n=5-6) outgrowth compared to hydrogels cultured with PSG1-His (*p*=0.019). Fold change in outgrowth area did not differ on days 1 and 3 of culture; however, on day 2, fold change in outgrowth area was significantly higher in samples cultured with PSG1-Fc compared to samples cultured with PSG1-His (One-way ANOVA, n=5-6, *p*=0.0078). Although we did not observe differences in cell viability between groups on day 3; however, cytotoxicity was found to be increased (One-way ANOVA, n=5-6) in PSG1-Fc samples compared to control (*p*=0.0082) and Fc-control samples (*p*=0.0071). Taken together, these results suggest that PSG1-Fc increases trophoblast motility.

**Figure 4.**
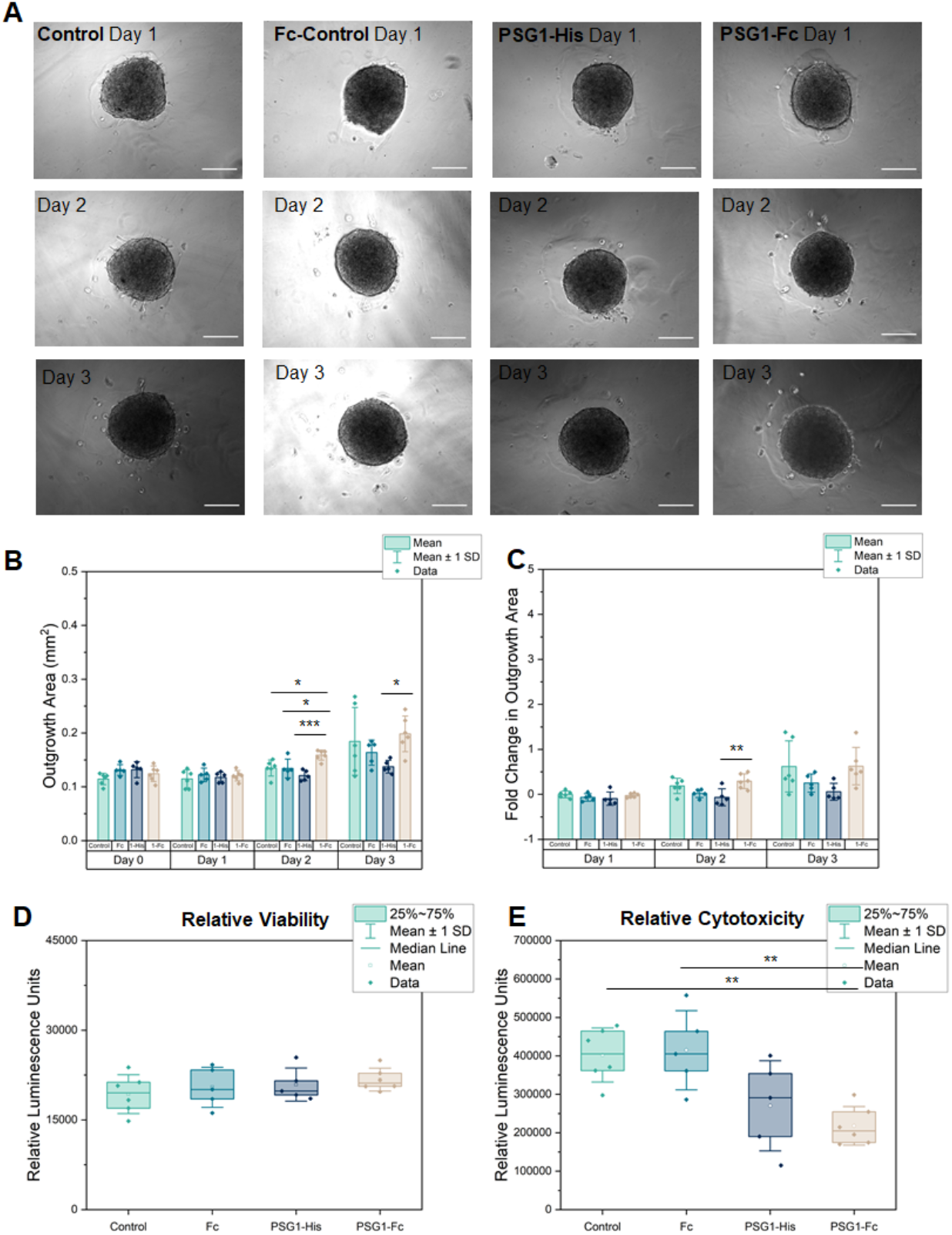
Invasion Pattern Analysis After the Addition of PSG1-Fc and PSG1-His to Swan71 Spheroids Encapsulated in Methacrylamide-Functionalized Gelatin (GelMA) Hydrogels. (A) Representative bright field images of 4,000 cell spheroids encapsulated in GelMA hydrogels from day 1 to day 3 for Control samples, Fc-Control samples (60 μg/ml), PSG1-His samples (60 μg/ml), and PSG1-Fc samples (60 μg/ml). Scale bar, 200 μm. (B) Average spheroid outgrowth area in GelMA hydrogels over 3 days. Data are displayed as individual data points. Bar represents mean ± standard deviation. Data were analyzed on each day between groups (Control n=6 technical replicates, Fc-Control (Fc) n=5 technical replicates, PSG1-His (1-His) n=5 technical replicates, and PSG1-Fc (1-Fc) n=6 technical replicates). Day 0, 1: One-way ANOVA. Day 2: One-way ANOVA with Tukey post hoc analysis, * p < 0.05, *** p < 0.001. Day 3: Welch’s ANOVA with Games-Howell post hoc analysis, * p < 0.05. (C) Spheroid outgrowth area fold change compared to initial area (day 0) over 3 days. Data are displayed as individual data points. Bar represents mean ± standard deviation. Data were analyzed on each day between groups (Control n=6 technical replicates, Fc-Control (Fc) n=5 technical replicates, PSG1-His (1-His) n=5 technical replicates, and PSG1-Fc (1-Fc) n=6 technical replicates). Day 1: Kruskal-Wallis ANOVA. Day 2: One-way ANOVA with Tukey post hoc analysis, ** p < 0.01. Day 3: One-way ANOVA. (D) Relative viability of encapsulated spheroids at day 3 from CellTiter-Glo® 3D Viability Assay. Data are presented as individual data points overlaying box plots with the median denoted by a line, mean denoted by a square, and whiskers represent the mean ± standard deviation. One-way ANOVA. Control n=6 technical replicates, Fc-Control (Fc) n=5 technical replicates, PSG1-His n=5 technical replicates, and PSG1-Fc n=6 technical replicates. (E) Relative cytotoxicity of encapsulated spheroids at day 3 from measured from lactate dehydrogenase release via LDH-Glo® Cytotoxicity Assay. Data are presented as individual data points overlaying box plots with the median denoted by a line, mean denoted by a square, and whiskers represent the mean ± standard deviation. One-way ANOVA with Tukey post hoc analysis. Control n=6 technical replicates, Fc-Control (Fc) n=5 technical replicates, PSG1-His n=5 technical replicates, and PSG1-Fc n=6 technical replicates, ** p < 0.01.

### 2.5. Swan71 Trophoblast Spheroids Are Capable of Matrix Deposition in 3D

Cell motility can be strongly influenced by dynamic remodeling of the extracellular matrix. We investigated bulk nascent protein production of encapsulated Swan71 trophoblast spheroids for days 1 through 3 of culture via a recently described metabolic labeling approach (utilizing cell media supplemented with the methionine-analog AHA) (Loebel et al., 2020). We observed significant nascent protein production as well as accumulation in close proximity to trophoblast cells in three-dimensional hydrogels (Figure 5). These observations indicate that nascent protein deposition occurs relatively rapidly within hydrogels and likely occurs throughout the entirety of these experiments (3 days).

**Figure 5.**
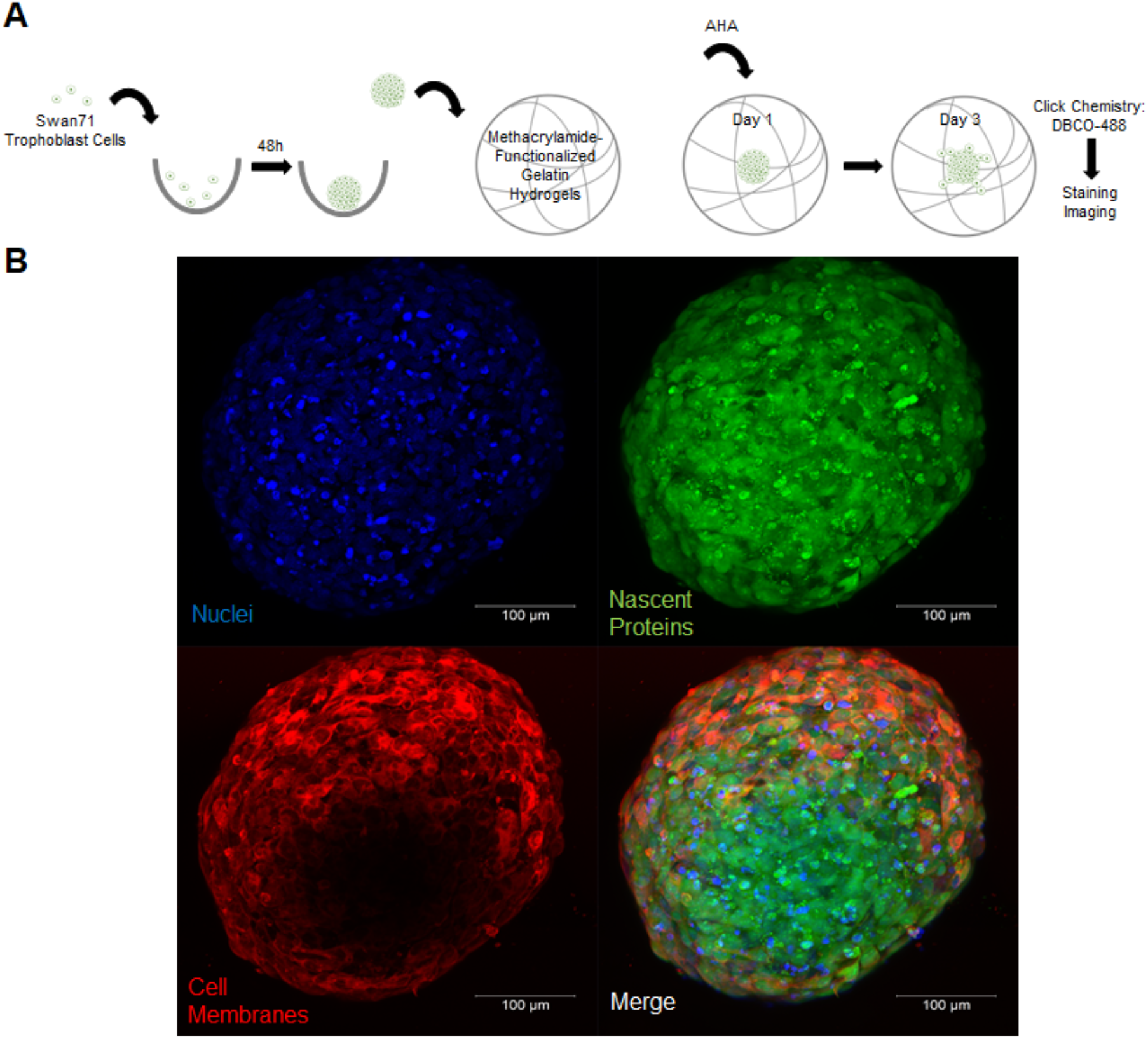
Matrix Remodeling by Encapsulated Swan71 Spheroids. (A) Schematic of experimental procedure. The methionine analog azidohomoalanine (AHA) was added to spheroids on day 1 of culture and replaced daily. On day 3, a click chemistry reaction was performed using DBCO-488, a fluorophore-conjugated cyclooctyne, to visualize nascent proteins. Staining was performed after the click chemistry reaction to visualize cells. (B) Representative maximum intensity projection of encapsulated Swan71 spheroid on day 3 of culture stained for bulk nascent protein production

## 3. Discussion

Many mechanisms driving early pregnancy and implantation remain unknown and poorly understood. Because of significant differences between animal and human pregnancy as well as the ethical concerns regarding studying early pregnancy in humans, *in vitro* models are often used to study mechanisms of human reproduction. Multidimensional *in vitro* models offer the opportunity to examine cellular mechanisms of early pregnancy using human-derived cells in controlled, tunable microenvironments. The 3D nature of these models can be tuned to recapitulate biophysical properties from the native tissue microenvironment such as architecture, stiffness, and inclusion of heterogeneous cohorts of cells, each offering increased biofidelity compared to existing 2D models. Recent 3D models of trophoblast migration have used biomaterials as a matrix around invading trophoblast cells as a means of replicating trophoblast invasion through tissue (Buck et al., 2015; Chang et al., 2018; Kuo et al., 2016; Kuo et al., 2018; Kuo et al., 2019; Wang et al., 2013; Wang et al., 2012); however, many of these models often employ biomaterials such as Matrigel that have limitations. For example, Matrigel has been found to have significant batch-to-batch variations in biomechanical properties, extracellular matrix content, and protein content (Aisenbrey and Murphy, 2020; Hughes et al., 2010; Vukicevic et al., 1992). The GelMA hydrogel system used in this work offers a number of potential advantages specific for investigating trophoblast activity. The gelatin macromer used to create the GelMA hydrogel contains RGD cell binding motifs and MMP sites which allows for cell binding and matrix remodeling (Loessner et al., 2016; Schuurman et al., 2013; Yue et al., 2015). The use of photoinitiator allows for photopolymerization of GelMA under UV light, which covalently crosslinks the hydrogels and renders the hydrogels relatively homogenous in structure (Loessner et al., 2016; Mahadik et al., 2017; Mahadik et al., 2015; Schuurman et al., 2013; Yue et al., 2015). The biophysical properties of GelMA, including stiffness and crosslinking density, can be tuned by manipulating the degree of methacrylamide functionalization along the gelatin macromer during GelMA synthesis or via crosslinking intensity (Gilchrist et al., 2019b; Loessner et al., 2016; Schuurman et al., 2013; Yue et al., 2015; Zambuto et al., 2019b). This makes the GelMA system amenable to creating libraries of monolithic environments or spatially-graded hydrogels containing a linear gradient of embossed stiffness or matrix-bound proteins to locally influence cell behavior (Gilchrist et al., 2019b; Mahadik et al., 2015; Pedron et al., 2017). Other groups have used microfabrication techniques such as photopatterning, micromolding, self-assembly, and 3D printing to further customize the three-dimensional environment (Schuurman et al., 2013; Yue et al., 2015). Although this study used unmodified GelMA hydrogels this versatility of the GelMA system provides exciting opportunities to examine trophoblast motility in increasingly complex, bioengineered systems.

We report Swan71 trophoblast motility in response to two different members of the human PSG family. We demonstrate reproducible, consistent methodology, as indicated by similar values in outgrowth area, fold change in outgrowth area, and viability for control samples across multiple experiments. We also quantify cytotoxicity of spheroids; however, this assay system has limitations compared to the viability assay system. We observed variability across experiments for cytotoxicity measurements even though the viability was similar, which has previously been found to be a limitation for the LDH assay and should be addressed in future studies by reducing serum concentrations and adjusting the signal-to-noise ratio for the assay (Bopp and Lettieri, 2008). To validate our motility assay, we first examined trophoblast motility in response to a known promoter (EGF) and inhibitor (Nodal) of trophoblast invasion. Exogenously added EGF markedly increased trophoblast outgrowth area compared to control and Nodal samples and increased viability compared to control samples, consistent with previous reports in the literature that concentrate on the mechanistic role of EGF in trophoblast activity (Kuo et al., 2016; Staun-Ram et al., 2004; Zambuto et al., 2020). Interestingly, samples cultured in the presence of Nodal showed no difference in outgrowth area and viability compared to control samples. Nadeem et al. previously demonstrated that Nodal reduced trophoblast invasion in wound-healing assays, transwell assays, and first-trimester placental explant invasion when Nodal was overexpressed or added to culture media (Nadeem et al., 2011).

PSGs are some of the most abundant circulating trophoblastic proteins in maternal blood during human pregnancy (Moore and Dveksler, 2014; Rattila et al., 2019; Sorensen, 1984; Wurz et al., 1981); however, much remains unknown regarding their role in early implantation and pregnancy disorders. Due to lack of specific antibodies that can distinguish the individual PSGs, temporal and expression levels have not been determined at the protein level; however, studies evaluating PSG expression at the mRNA level in the first and third trimesters indicated that PSG1 is expressed at much higher levels than PSG9, a finding confirmed by a different laboratory when examining expression of PSGs in villous cytotrophoblasts (Camolotto et al., 2010; Shanley et al., 2013).

The various PSGs that have been examined so far share identical functions; therefore, the expansion and rapid evolution of the human PSG family has been explained by selection for increased dosage of PSG proteins, rather than for diversification of function. Both PSG1 and PSG9 activate latent transforming growth factor β (TGF-β, bind to integrins αIIb3, and α5β1, induce endothelial tube formation by binding to heparan sulfate proteoglycans and also bind to galectin-1 in a glycan-dependent manner (Jones et al., 2016; Rattila et al., 2019; Shanley et al., 2013; Warren et al., 2018). Interestingly, although lower than normal concentrations of PSG1 were reported in preeclampsia using a PSG1-specific enzyme-linked immunosorbent assay (ELISA), Blankley and co-workers, using mass spectrometry, reported that PSG9 and PSG5-derived peptides were increased in plasma at 15 weeks gestation in women suffering from early-onset preeclampsia when compared to women with healthy pregnancies (Blankley et al., 2013; Rattila et al., 2019). Although additional studies are required to confirm the potential differences in PSG1 and PSG9 expression and its association with pregnancy pathologies, we studied the effect of exogenously added PSG1 and PSG9 in trophoblast motility with this newly developed protocol.

To this end, we cultured encapsulated Swan71 spheroids with PSG9-Fc and quantified outgrowth area, viability, and cytotoxicity after 3 days of culture. We found that compared to control and Fc-control samples, PSG9-Fc reduced the trophoblast outgrowth area. This study suggests that PSG9-Fc reduces trophoblast invasion; however, in order to fully understand the mechanism, additional studies must be performed. We subsequently examined the influence of exogenous PSG1-Fc and PSG1-His on trophoblast motility. Previous studies showed staining of PSG1 in extravillous trophoblasts in distal placental areas, suggesting that there is likely increased expression of this PSG and potentially one or more of the other PSGs recognized by the monoclonal antibody utilized for the studies (PSG6, PSG7, and PSG8) in extravillous trophoblasts with an invasive phenotype (Rattila et al., 2019). However, previous studies using two-dimensional wound healing assays with Swan71 or transwell invasion assays using the HTR-8 SVneo extravillous trophoblast cell line did not identify shifts in cell invasion in response to soluble PSG1 (Rattila et al., 2019). We employed our trophoblast motility assay to investigate the influence of two variants of PSG1 (PSG1-Fc and PSG1-His). These two proteins differ from each other by the tag fused to the PSG1 sequence, which results in a dimeric protein in the case of PSG1-Fc, and in the length of the carboxy-terminal end following the B2 domain (1a slice variant and 1d slice variant, respectively). Interestingly, we observed an increase in outgrowth area between control samples and samples incubated with PSG1-Fc on day 2 and day 3 of culture. Although we did not observe differences in outgrowth area between control samples and PSG1-His, we observed reduced cytotoxicity in response to PSG1-His versus control. We expected to observe similar results with PSG1-Fc and PSG1-His based on our prior results analyzing PSG1 binding to their binding partners; however, PSG1-His seems to cause a downward reduction in outgrowth area compared to control samples. Nevertheless, it is possible that PSG1-Fc and PSG1-His could have the same effects on trophoblast invasion and cell viability, but a limitation of these studies is that all proteins were utilized at a single concentration. Dose-response studies may provide additional insights into the differences in the activity of the proteins in the spheroid outgrowth assay. In addition, although we observed very low expression of PSG1 in the supernatant of Swan71 cells by ELISA (approximately 525 pg/mL secreted over a 48 h period after seeding 100,000 cells in a 6 well plate); transfection of the Swan 71 cells with the PSG1 and PSG9 rather than addition of the proteins to the media may provide additional valuable information on their activity as regulators of trophoblast motility.

Finally, we demonstrate the three-dimensional trophoblast motility assay also provides an avenue to explore trophoblast mediated matrix remodeling. Remodeling is a powerful force central to successful trophoblast invasion. We have previously used the GelMA hydrogel as a well-characterized platform to examine processes of cell mediated degradative and biosynthetic remodeling via assessments of stiffness, gene expression, and secreted biomolecules (Gilchrist et al., 2019a). Here, we adapted a recently described metabolic labeling approach to examine qualitative nascent protein production by Swan71 spheroids. We stained cells for bulk nascent protein production after 3 days of culture. Spheroids of Swan 71 trophoblasts secrete proteins within the hydrogels in sufficient quantities that their local accumulation can be visualized. Matrix remodeling plays a key role in trophoblast invasion so methods such as these will enable researchers to probe questions relating to the role of the extracellular matrix on trophoblast cell migration (Cha et al., 2012; Cohen and Bischof, 2007; Knofler, 2010; Norwitz et al., 2001). Future work will determine the specific proteins secreted by the cells to determine if matrix deposition plays a role on trophoblast motility.

In conclusion, multidimensional biomaterial platforms offer the opportunity to provide 3D tissue microenvironments to perform mechanistic studies to elucidate physiological and pathophysiological processes challenging to study *in vivo*. Our approach to recapitulate aspects of the endometrial microenvironment (e.g., tissue stiffness, spheroid culture, migration through a matrix) to probe questions relating to trophoblast invasion employs spheroids encapsulated in GelMA hydrogels to perform quantitative motility and viability assays. We used these assays to determine trophoblast response to soluble PSG9 and PSG1. Our results show that a spheroid-based motility has the sensitivity to rapidly (within 3 days) quantify differences in trophoblast invasion. Notably, we find PSG9-Fc reduces trophoblast motility while PSG1-Fc increases trophoblast motility compared to control samples. And further, these cells secrete significant new protein content that can be readily visualized. Taken together, this 3D trophoblast motility model has the potential to allow us to better understand trophoblast interactions with the extracellular matrix through which they are migrating so we can develop a deeper understanding of the mechanisms associated with early implantation.

## 4. Experimental Procedures

### 4.1. Methacrylamide-Functionalized Gelatin (GelMA) Synthesis and Hydrogel Fabrication

Methacrylamide-functionalized gelatin was synthesized as described previously using a method developed by Shirahama et al. (Shirahama et al., 2016; Zambuto et al., 2019b, 2020). GelMA was determined to have a 57% degree of functionalization as determined by ^1^H-NMR and a stiffness of approximately 2 kPa as previously determined by compression testing (Zambuto et al., 2020). 5 wt% polymer solutions were created by dissolving lyophilized GelMA in phosphate buffered saline (PBS; Lonza, 17-516F) at 37°C, adding 0.1% w/v lithium acylphosphinate (LAP) as photoinitiator, and polymerizing hydrogels under UV light for 30 seconds (λ=365 nm, 7.14 mW cm^−2^; AccuCure Spot System ULM-3-365). Each hydrogel was fabricated from 20 μL prepolymer solution pipetted into custom circular Teflon molds (5 mm diameter, 1 mm height).

### 4.2 Generation of Recombinant PSG1 and PSG9 and Fc Control Proteins

PSG1-His was harvested from the supernatant of a stably transfected CHO-K1 single-cell clone established in our laboratory and grown in a hollow fiber cartridge (FiberCell Systems, MD, USA) as previously described (Ballesteros et al., 2015). The Fc fusion proteins were generated by cloning the PSG1a cDNA (UniProt KB-P11464) and PSG9 cDNA (NM_002784) into the pFuse-IgG1e3-Fc1 vector (InvivoGen, San Diego, CA). The Fc control protein was derived from the pFuse-IgG1e3-Fc2 (InvivoGen). For the generation of full length PSG1-Fc and PSG9-Fc, the plasmids were transfected into ExpiCHO cells (Thermo Fisher Scientific, Waltham, MA, USA) following the manufacturer’s recommendations and the supernatants were collected 6 days post-transfection. Proteins were purified from the clarified supernatants on an anti-PSG1 mAb#4 column (for PSG1-His) and protein A columns (for PSG1-Fc, PSG9-Fc, Fc control) in an AKTA pure chromatography system (Cytiva, Marlborough, MA). All proteins were eluted with 0.1 M glycine buffer pH 2.7, followed by immediate neutralization with 1 M Tris-HCl pH 8 (Thermo Fisher Scientific). Following elution from the column, the fractions containing the proteins were pooled, concentrated, and buffer-exchanged with phosphate-buffered saline (PBS) using an Amicon Ultra centrifugal filter unit with a 10 kDa cutoff membrane (Millipore, Burlington, MA). To determine the protein concentration, the purified proteins were separated on 4–12% NuPAGE Bis-Tris gels (Thermo Fisher Scientific) at different dilutions alongside known concentrations of BSA used as standards. The gels were stained with GelCode Blue (Thermo Fisher Scientific), and the proteins were quantitated by densitometry using a BioRad Gel Doc EZ Imager.

### 4.3. Swan71 Spheroid Motility Assays

#### 4.3.1. Swan71 Cell Culture and Maintenance

The telomerase immortalized first trimester trophoblast cell line Swan71 was provided by Dr. Gabriela Dveksler. Cells were maintained in DMEM (Gibco, 11995-065) supplemented with 10% fetal bovine serum (FBS; R&D Systems, S11150H), 1% penicillin/streptomycin (Thermo Fisher, 15140122), and 500 ng/mL puromycin (Sigma-Aldrich, P8833). Medium was replaced every other day of culture. All cultures were grown in a humidified incubator at 37°C with 5% CO_2_. Cells were routinely tested for mycoplasma every 6 months to ensure cell quality (MycoAlert™ Mycoplasma Detection Kit; Lonza).

#### 4.3.2. Spheroid Motility Assays

Spheroid motility assays were performed as previously described by our group (Zambuto et al., 2019b, 2020). Spheroids and encapsulated spheroids were cultured in DMEM supplemented with 2% FBS, and 1% penicillin/streptomycin. Briefly, 4,000 Swan71 cells were added to each well of a round bottom plate (Corning, 4515). The plate was placed on a shaker at 60 rpm in a humidified incubator at 37°C with 5% CO_2_ for 48 hours to allow spheroids to form. After 48 hours, individual spheroids were pipetted onto Teflon molds and 20 μL prepolymer solution was added to each mold well. Each spheroid was gently moved to the center of each hydrogel using a pipette tip. Hydrogels were then polymerized and added to 48 well plates containing 500 μL medium per well. Once all samples were fabricated, each hydrogel was imaged and then medium was replaced with 500 μL medium supplemented with relevant biomolecules.

Recombinant human epidermal growth factor (EGF, 5 ng/mL; Sigma-Aldrich, E9644) was used as an invasion promoter and recombinant human nodal (250 ng/mL; R&D Systems, 3218-ND-025/CF) was used as an invasion inhibitor. Additional control samples were cultured in medium without biomolecules. Encapsulated spheroids were imaged daily using a Leica DMI 4000 B microscope (Leica Microsystems). Total outgrowth area was calculated using the measure tool in Fiji by manually tracing spheroids three times and taking the average of these three measurements. Fold change was calculated for each hydrogel from average outgrowth area by comparing average outgrowth area on days 1, 2, and 3 to average area on day 0 (day of encapsulation) using Eq. 1.

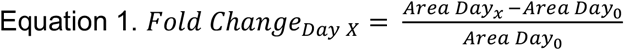

#### 4.3.3 CellTiter-Glo® 3D Viability Assay

Viability was quantified using the CellTiter-Glo® 3D Viability Assay (Promega) as described previously using the manufacturer’s instructions (Zambuto et al., 2020). Briefly, samples and reagents were equilibrated to room temperature for at least 30 minutes prior to completing the assay. Medium was removed from each hydrogel and 400 μL of a 1:1 solution of CellTiter-Glo® and medium was added to each hydrogel. Samples were incubated for 1 hour at room temperature on a shaker. Following incubation, 100 μL triplicates for each sample were added to an opaque, white-walled plate, including a blank consisting of only the stock solution. Plate luminescence was read immediately using a plate reader (BioTek Synergy HT Plate Reader and Gen5 Software, BioTek Instruments, Inc.). Relative luminescence for each sample was calculated by subtracting the average blank luminescence value from the average luminescence value for each sample.

#### 4.3.4. LDH-Glo™ Cytotoxicity Assay

Cytotoxicity was quantified using the LDH-Glo™ (Promega, J2380) as described by the manufacturer’s instructions. Briefly, samples were prepared by combining 4 μL of medium per hydrogel and diluting it in 96 μL LDH (lactate dehydrogenase) storage buffer (200 mM Tris-HCl (Roche, 10812846001), 10% glycerol (Promega, H5433), 1% bovine serum albumin (BSA; Sigma-Aldrich, A4503); filter sterilized and stored at 4°C). This dilution was calculated to fall within the linear regime of the assay (data not shown). Samples were used immediately or stored at −80°C until use. Prior to running the assay, all reagents were equilibrated to room temperature. Duplicate wells were prepared for each hydrogel by adding 50 μL of sample and 50 μL prepared LDH Detection Reagent per well in 96-well opaque, white-walled plates. Samples were protected from light and incubated at room temperature for 60 minutes. After 60 minutes, luminescence was read immediately using a plate reader with a 1 second integration time (BioTek Synergy HT Plate Reader and Gen5 Software, BioTek Instruments, Inc.). Relative luminescence for each sample was calculated by subtracting the luminescence from the sample blank (cell medium) from each hydrogel sample.

### 4.4. Spheroid Immunofluorescent Staining and Imaging

Encapsulated spheroids were stained as described previously (Zambuto et al., 2020). On days 1 and 3 of culture, encapsulated spheroids were fixed in 4% formaldehyde in PBS for 15 minutes followed by three PBS washes. All subsequent steps were performed on a shaker at room temperature unless otherwise noted. Samples were permeabilized for 15 minutes in a 0.1% Tween 20 (Fisher Scientific, BP337) in PBS. Following permeabilization, 300 μL of working solution was added to each sample (1 μL Phalloidin-iFluor 488 Reagent (Abcam, ab176753) per 1 mL 1% BSA in PBS) for 90 minutes and samples were protected from light. After incubation, samples were washed in PBS 4×20 minutes. Samples were then stained with Hoechst (1:2000 in PBS; ThermoFisher, H3570) for 30 minutes followed by one PBS wash. Samples were stored at 4°C in PBS until imaged. Samples were imaged using a Zeiss LSM 710 Confocal Microscope and 20X objective. Two samples were imaged per day and one Z-stack was taken per sample. Maximum intensity projection images were generated using ZEN (black edition; Zeiss).

### 4.5. Nascent Protein Staining and Imaging

Nascent protein staining was performed as previously described by Loebel et al. (Dieterich et al., 2007; Loebel et al., 2020; Loebel et al., 2019). Starting on day 1 of culture, encapsulated spheroids were incubated with the methionine analog azidohomoalanine (Click-iT AHA; Invitrogen, C10102; 100 μM) in methionine-free DMEM (custom made; School of Chemical Sciences Cell Media Facility, University of Illinois at Urbana-Champaign) supplemented with 1% penicillin/streptomycin and 2% FBS. AHA was replenished daily. On day 3 of culture, live cells were washed two times with sterile PBS, incubated for 40 minutes in AFDye 488 DBCO (Click Chemistry Tools, 1278-1; 30 μM) at 37°C, washed three times with sterile PBS, and then fixed in 4% formaldehyde in PBS for 15 minutes followed by three PBS washes. Samples were stored at 4°C in PBS until stained. Samples were incubated in CellMask™ Deep Red Plasma Membrane Stain (1:1000; Invitrogen, C10046) for 40 minutes at 37°C followed by three PBS washes. Samples were then incubated in Hoechst at room temperature for 30 minutes (1:2000 dilution in PBS) followed by one PBS wash. Samples were stored at 4°C in PBS until imaged. Samples (n=3) were imaged using a Zeiss LSM 710 Confocal Microscope and 20X objective. One Z-stack was taken per sample and maximum intensity projection images were generated using ZEN (black edition; Zeiss).

### 4.6. Statistics

OriginPro 2020 (Origin Lab) and RStudio were used for statistical analysis. Underlying assumptions for each statistical test were tested prior to analyzing data. Normality was determined using the Shapiro-Wilkes test and homoscedasticity was determined using Levene’s test. For each experiment, n=5-8 independent spheroids were analyzed per sample group. Normal, homoscedastic data were analyzed using a one-way analysis of variance (ANOVA) and post hoc Tukey Test. Normal, heteroscedastic data were analyzed using Welch’s ANOVA and post hoc Games-Howell Test. Non-normal, homoscedastic data were analyzed using Kruskal-Wallis ANOVA and post hoc Dunn’s Test. Non-normal, heteroscedastic data were analyzed using Welch’s Heteroscedastic F Test with Trimmed Means and Winsorized Variances and post hoc Games-Howell Test. Significance was defined as *p*<0.05. Data are presented as mean ± standard deviation unless otherwise noted.

## Acknowledgements

Research reported was supported by the National Institutes of Diabetes and Digestive and Kidney Diseases of the National Institutes of Health under Award Numbers R01 DK0099528 (B.A.C.H.), National Institute of Allergy and Infectious Diseases under Award R21 AI1290918 (G.D) and by the National Institute of Biomedical Imaging and Bioengineering of the National Institutes of Health under Award Numbers R21 EB018481 (B.A.C.H.) and T32 EB019944 (S.G.Z.). The content herein is solely the responsibility of the authors and does not necessarily represent the official views of the National Institutes of Health or the Department of Defense. The authors also gratefully acknowledge additional funding provided by the Department of Chemical & Biomolecular Engineering and the Carl R. Woese Institute for Genomic Biology at the University of Illinois at Urbana-Champaign. The authors thank Dr. Gil Mor (Yale University School of Medicine, New Haven, CT) for providing the Swan71 cells. The authors also thank the School of Chemical Sciences Cell Media Facility (Dr. Sandy McMasters) at the University of Illinois at Urbana-Champaign for assistance with cell media for the nascent protein experiment and the Institute for Genomic Biology Core Facilities (Dr. Austin Cyphersmith) at the University of Illinois at Urbana-Champaign for assistance with confocal imaging.

## Author Contributions

S.G.Z. designed and conducted experiments, analyzed resultant data, and wrote the manuscript. S. R. generated the recombinant PSGs. G.D. conceived of the study and edited the manuscript. B.A.C.H. provided experimental guidance and edited the manuscript.

